# Scalable variational inference for low-rank spatio-temporal receptive fields

**DOI:** 10.1101/2022.08.12.503812

**Authors:** Lea Duncker, Kiersten M. Ruda, Greg D. Field, Jonathan W. Pillow

## Abstract

An important problem in systems neuroscience is to characterize how a neuron integrates sensory inputs across space and time. The linear receptive field provides a mathematical characterization of this weighting function, and is commonly used to quantify neural response properties and classify cell types. However, estimating receptive fields is difficult in settings with limited data and correlated or high-dimensional stimuli. To overcome these difficulties, we propose a hierarchical model designed to flexibly parameterize low-rank receptive fields. The model includes Gaussian process priors over spatial and temporal components of the receptive field, encouraging smoothness in space and time. We also propose a new temporal prior called temporal relevance determination (TRD), which imposes a variable degree of smoothness as a function of time lag. We derive a scalable algorithm for variational Bayesian inference for both spatial and temporal receptive field components and hyperparameters. The resulting estimator scales to high-dimensional settings in which full-rank maximum likelihood or *a posteriori* estimates are intractable. We evaluate our approach on neural data from rat retina and primate cortex, and show that it substantially out-performs a variety of existing estimators. Our modeling approach will have useful extensions to a variety of other high-dimensional inference problems with smooth or low-rank structure.

## 1 Introduction

A key problem in computational and systems neuroscience is to understand the information carried by neurons in the sensory pathways [1, 2]. A common approach to this problem is to estimate the linear receptive field (RF), which provides a simple characterization of a neuron’s stimulus-response properties. The RF consists of a set of linear weights that describe how a neuron integrates a sensory stimulus over space and time [1, 3–11]. Estimating the RF from imaging or electrophysiological recordings can thus be seen as a straightforward regression problem. However, characterizing RFs in realistic settings poses a number of important challenges.

One major challenge for RF estimation is the high dimensionality of the inference problem. The number of RF coefficients is equal to the product of the number of spatial stimulus elements (e.g. pixels in an image) and number of time lags in the temporal filter. In realistic settings this can easily surpass thousands of coefficients. Classic RF estimators such as the least-squares regression suffer from high computational cost and low statistical power in high-dimensional settings. The computational cost of the regression estimate scales cubically with the number of RF coefficients, while memory cost scales quadratically. Moreover, the standard regression estimate does not exist unless there are more samples than dimensions, and typically requires large amounts of data to achieve a high level of accuracy. High dimensionality is thus a limiting factor in terms of both computational resources and statistical power.

Past efforts to improve statistical power, and thereby reduce data requirements, have relied on various forms of regularization. Regularization reduces the number of degrees of freedom in the RF by biasing the estimator towards solutions that are more likely *a priori*, and can thus provide for better estimates from less data. Common forms of regularizaton involve smoothness or sparsity assumptions, and have been shown to outperform maximum likelihood (ML) estimation in settings of limited data or correlated stimuli [12–18]. However, the computational cost of such estimators is generally no better than that of classical estimators, and may be worse due to the need to optimize hyperparameters governing the strength of regularization. These poor scaling properties make it difficult to apply such estimators to settings involving high-dimensional stimulus ensembles.

In this paper, we propose to overcome these difficulties using a model that parametrizes the RF as smooth and low rank. A low rank receptive field can be described as a sum of a small number of space-time separable filters [13, 19–22]. This choice of parametrization significantly reduces the number of parameters in the model and is the key to scalability of our method. To achieve smoothness, we use Gaussian process (GP) priors over both the spatial and temporal filters composing the low-rank RF. For the temporal filters, we introduce the temporal relevance determination (TRD) prior, which uses a novel covariance function that allows for increasing smoothness as a function of time lag. To fit the model, we develop a scalable algorithm for variational Bayesian inference for both receptive field components and hyperparameters governing shrinkage and smoothness. The resulting estimator achieves excellent performance in settings with correlated stimuli, and scales to high-dimensional settings in which full-rank estimators are intractable.

The paper is organized as follows. Sections 2-3 provide a review of relevant background and existing literature: Section 2 introduces the linear encoding model and low rank linear receptive fields; Section 3 reviews previously proposed priors for receptive fields. In Section 4, we introduce temporal relevance determination (TDR), a new prior for temporal receptive fields, while in section 5 we introduce our variational low-rank (VLR) receptive field estimator. Section 6 shows applications to simulated as well as real neural datasets, and, at last, Section 7 provides discussion of our results and suggests directions for future work.

## 2 The Linear-Gaussian encoding model

A classic approach to neural encoding is to formulate a parametric statistical model that describes the mapping from stimulus input to neural response [1, 2, 17, 23, 24]. Here we focus on linear models, where the neural response is described as a linear or affine function of the stimulus plus Gaussian noise [17, 25] (see Fig. 1). Formally, the model can be written:

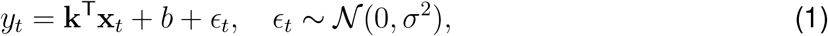

where *y*_*t*_ is the neuron’s scalar response at the *t*’th time bin, **k** is the vector receptive field, **x**_*t*_ is the vectorized stimulus at the *t*’th time bin, *b* is an additive constant or bias, and *ϵ*_*t*_ denotes zero-mean Gaussian noise with variance *σ*^2^.

**Figure 1:**
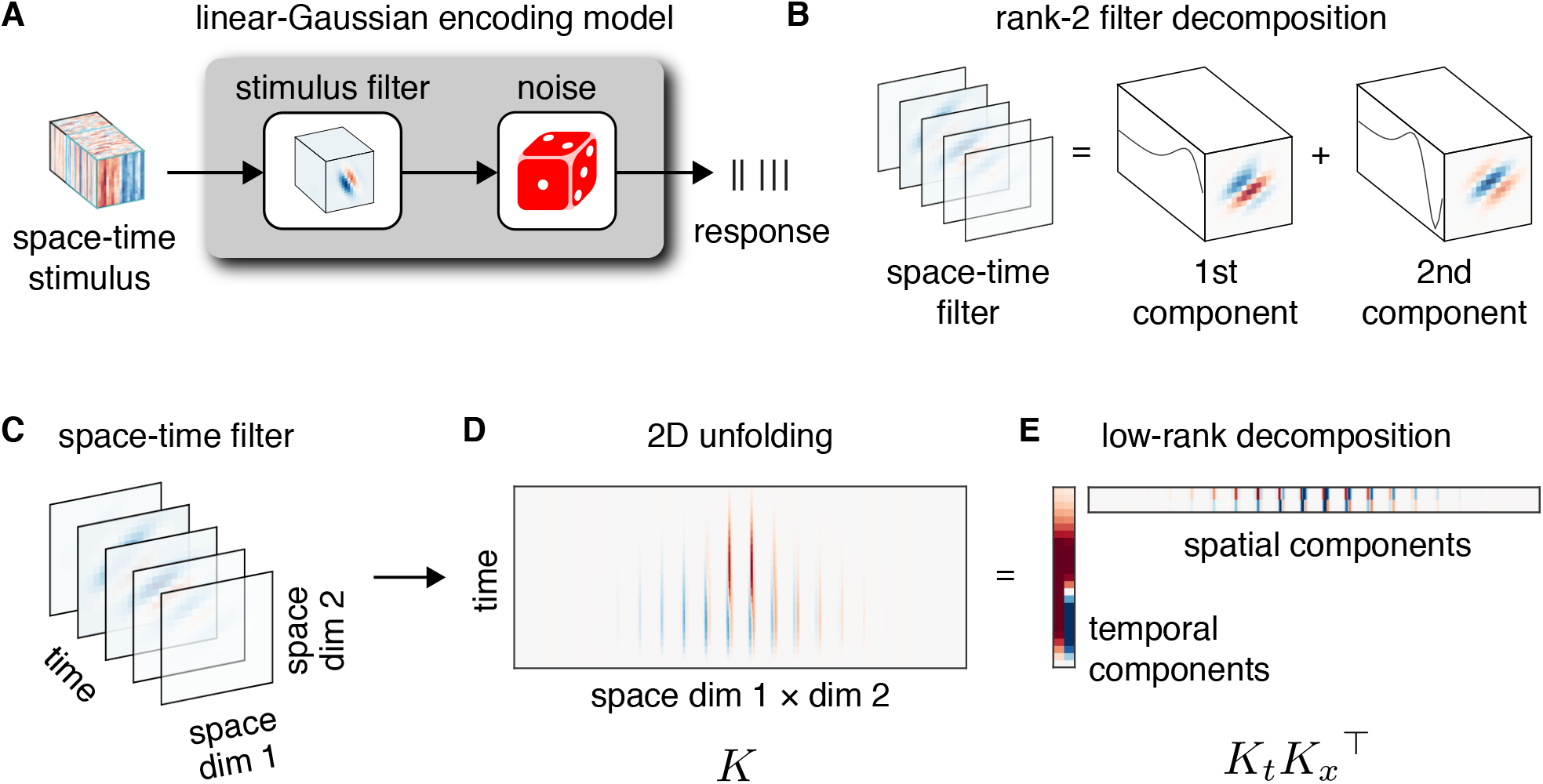
**A**: The linear-Gaussian encoding model is a 2-stage model, consisting of a linear stage followed by a noise stage. First, the high-dimensional spatio-temporal stimulus is filtered with the linear receptive field, which describes how the neuron integrates the stimulus values over space and time. The output of this filter is a scalar that gives the expected response at a single time bin. We then add noise to obtain the neural response for the given time bin. **B**: A low-rank spatio-temporal receptive field can be described as a sum of two space-time-separable (rank 1) filters. **C-D**: Illustration of different STRF representations as a third order tensor (**C**), matrix (**D**) or in terms of low-rank factors (**E**).

### 2.1 The receptive field tensor

The model description above neglects the fact that the stimulus and receptive field are often more naturally described as multi-dimensional tensors. In vision experiments, for example, the stimulus and receptive field are typically third order tensors, with two dimensions of space and one dimension of time. In this case, the full stimulus movie shown during an entire experiment can be represented by a tensor of size 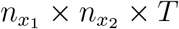, consisting of 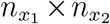 pixel images over *T* total time bins or frames. In this case, a given element **X**_*i,j,t*_ in the stimulus tensor could represent the light intensity in a grayscale image at time *t* at spatial location (*i, j*) in the image.In such settings, the spatio-temporal receptive field is also naturally defined as a tensor, 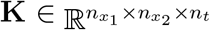, with weights that determine how the neuron integrates light over the 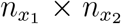 spatial pixels and the *n*_*t*_ preceding time bins (Figure 1C). Thus, the dot product between the receptive field **k** and vector stimulus **x**_*t*_ in (eq. 1) is equal to the following linear function defined by summing over the product of all corresponding elements of the RF tensor **K** and corresponding portion of the stimulus tensor **X** at time *t* :

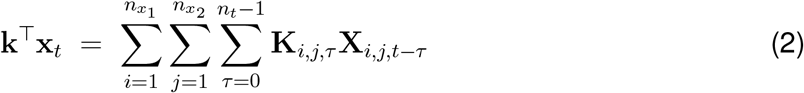

where **K** and **X**_*t*_ are both tensors of size 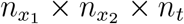. The vectorized RF and stimulus are thus given by **k** = vec(**K**), and **x**_*t*_ = vec(**X**_*i,j,t*−τ_) for 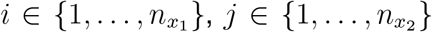 and *τ* − {1, …, *n*_*t*_}.

### 2.2 Low rank receptive fields

A key feature of neural receptive fields is that they can typically be described by a relatively small number of spatial and temporal components [13, 19–22, 26]. This means that we do not need to use a full set of 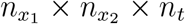 independent parameters to represent the coefficients in **K**. Instead, we can accurately describe **K** with a small number of spatial components, each corresponding to an image with 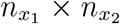 coefficients, and a corresponding number of temporal components, each with *n*_*t*_ coefficients. The number of paired spatial and temporal components needed to represent **K** is known as the rank of the tensor, which we denote *r*.

Figure 1B illustrates a scenario with *r* = 2, in which the tensor **K** is the sum of two rank-1 components, each defined as a single spatial and temporal weighting function. These rank-1 components are commonly referred to as “space-time separable” filters. Note that a rank *r* tensor has only 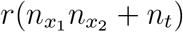 parameters, which generally represents a significant savings over the 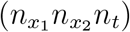 parameters needed to parametrize the full-rank tensor. Furthermore, having an explicit description of the temporal and spatial filters composing **K** increases interpretability of the RF.

For the purposes of modeling low-rank filters of this form, it is convenient to unfold the 3rd order receptive field tensor **K** into a matrix. Let 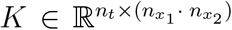 denote the matrix unfolding of the receptive field, where the two spatial dimensions have been concatenated (see Fig. 1C,D). This representation makes it possible to represent low-rank receptive fields with a product of matrices:

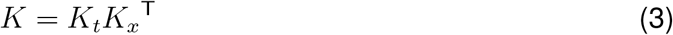

where 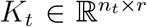 is a matrix whose columns are temporal filters, and 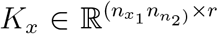 is a matrix whose columns are (reshaped) spatial filters (see Fig. 1E). In section 5 we will develop a Bayesian hierarchical model for efficient estimation of low-rank receptive fields using this parametrization.

## 3 Existing receptive field priors

In high-dimensional settings, or settings with highly correlated stimuli, receptive field estimation can be substantially improved by regularization. Here we review previously proposed prior distributions for regularizing receptive field estimates. The general family of priors that we consider takes the form of a zero-mean multivariate Gaussian distribution:

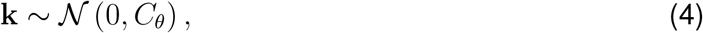

where *C*_*θ*_ denotes a covariance matrix that depends on a set of hyperparameters *θ*. Different choices of prior arise by selecting different functional forms for the covariance matrix *C*_*θ*_. We will review several popular choices of covariance below (see Fig. 2), before introducing a novel prior covariance in section 4.

**Figure 2:**
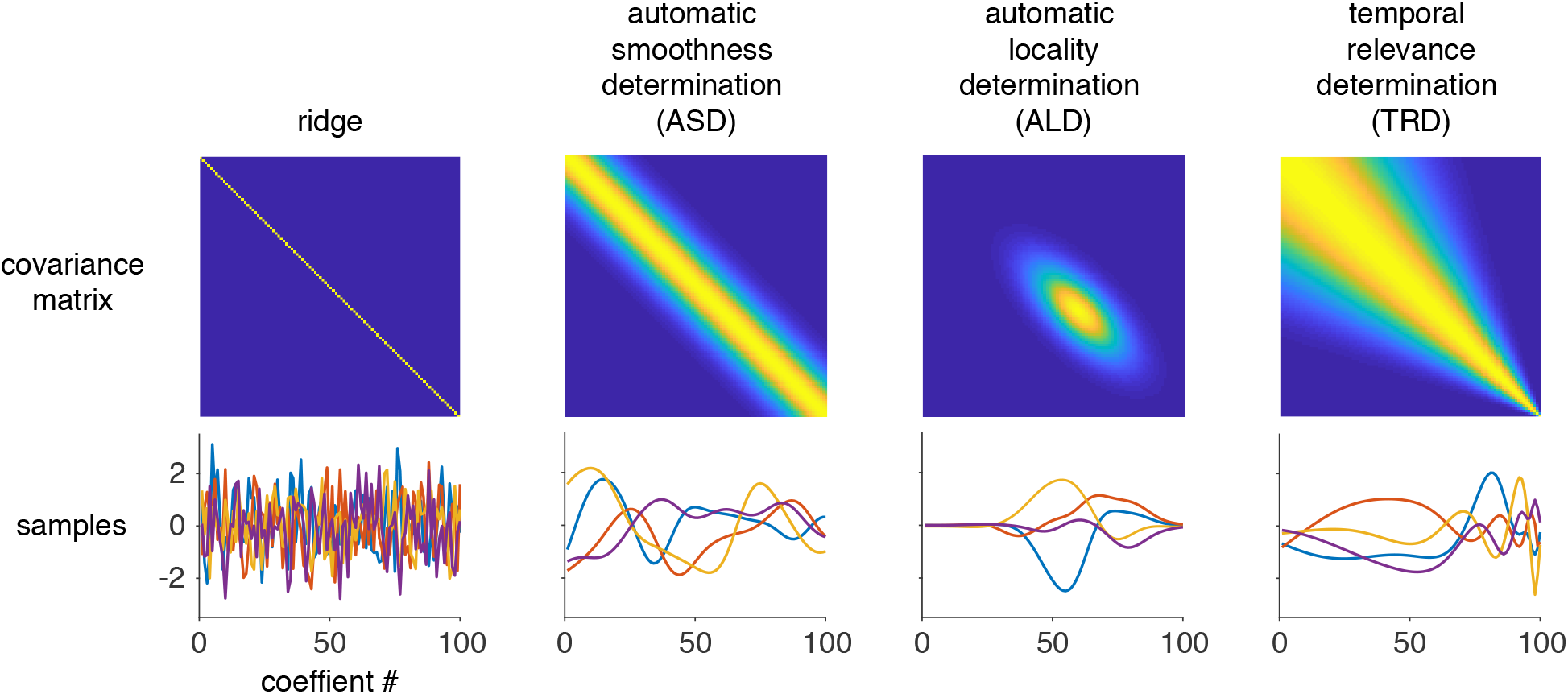
Illustration of different priors for use in receptive field estimation. Top row: prior covariance matrices Cθ under different choices of covariance function. Under the ridge prior, all off-diagonal elements of the covariance matrix are zero and receptive field coefficient are independent. In the ASD prior covariance, neighboring coefficients are highly correlated and the correlation decreases with increasing distance between coefficient locations in the RF. The ALD prior covariance contains high correlations for neighboring coefficients within a local region, and additionally sets the prior covariance of the RF coefficient to zero outside of this local region – a form of structured sparsity. Finally, in the TRD covariance matrix, the correlation of neighboring RF coefficients increases as a function of the coefficient location. Bottom row: Four samples from a multivariate Gaussian distribution with zero mean and covariance matrix shown above, illustrating the kinds of receptive fields that are typical under each choice of prior.

### 3.1 Ridge prior

Ridge regression [27] is classically viewed as an added *L*_2_ norm penalty on the receptive field weights in the context of least-squares regression. However, it also has a Bayesian interpretation as resulting from a zero-mean Gaussian prior with covariance given by

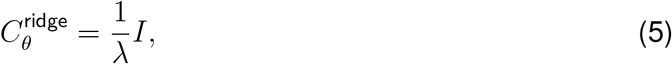

where the hyperparameter *θ* = {λ} is known as the ridge parameter, and *I* denotes the identity matrix. This prior has the effect of biasing the estimate towards zero, a phenomenon also known as “*L*_2_ shrinkage”.

### 3.2 Automatic Smoothness Determination (ASD)

The Automatic Smoothness Determination (ASD) prior [25] goes beyond shrinkage to incorporate the assumption that the receptive field changes smoothly in time and/or space. The ASD prior covariance matrix relies on the radial basis function (RBF) or “Gaussian” covariance function that is well-known in the Gaussian process literature [28]. The *i, j*’th element of this covariance matrix is given by:

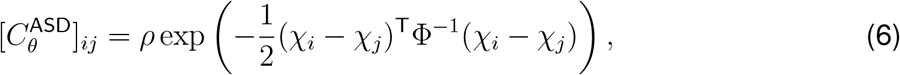

where {*χ*_*i*_} is a 3D vector containing the locations of RF pixels in space-time, thus indicating both the 2D spatial locations of the RF coefficients and the 1D temporal locations (or lags). And 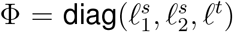 is a diagonal matrix containing “lengthscale” parameters. The covariance matrix is thus controlled by four hyperparameters, 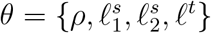. *ρ* is the marginal variance (and equivalent to 1/λ in the ridge prior above), and the lengthscale parameters 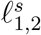 and *ℓ*^*t*^ determine the degree of smoothness in space and time, respectively. Recent work has exploited the Kronecker and Toeplitz structure of the ASD covariance matrix to show that it has an exact diagonal representation in the Fourier domain, which allows for dramatic improvements in computational efficiency [18].

### 3.3 Automatic Locality Determination (ALD)

The automatic Locality Determination (ALD) prior [17] goes beyond smoothness of the ASD prior by encouraging RF coefficients to be localized in space, time, and in spatio-temporal frequency.

It relies on a covariance function that encourages both the space-time coefficients and the spatio-temporal frequency components of the RF to be zero outside of some localized region, resulting in a form of “structured sparsity” [29]. This prior includes the ASD smoothness prior as a special case, namely when the region of non-zero spatio-temporal frequency components is centered at zero, and there is no locality in space or time. However, the ALD prior also allows for band-pass filters, in which the RF is composed primarily of intermediate frequencies.

The ALD covariance matrix can be written in terms of a pair of diagonal matrices a discrete Fourier transform matrix:

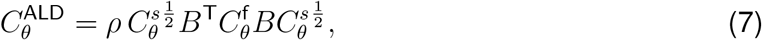

where 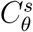 and 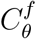 and are diagonal matrices that specify a region of nonzero coefficients in space-time and the Fourier domain, respectively, and *B* is the discrete-time Fourier transform matrix.

The space-time locality matrix is a diagonal matrix with diagonal elements given by

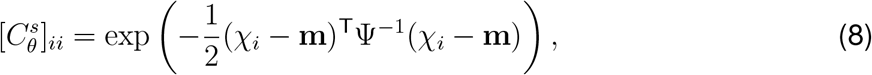

where {*χ*_*i*_} are the locations of RF pixels in space-time, and **m** and Ψ denote the mean and covariance of the region where the RF coefficients are non-zero. The Fourier-domain locality matrix takes a similar form:

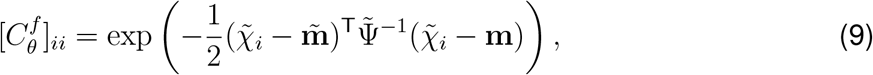

where 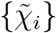 are the spatio-temporal frequencies for each Fourier coefficient, and 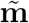 and 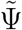 denote the mean and covariance of the region in Fourier space where RF Fourier coefficients are non-zero. The hyperparameters governing the ALD prior are thus 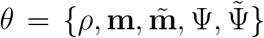. As before, *ρ* governs the marginal variance of the RF coefficients, analogous to the ridge parameter. For a 3D tensor receptive field, **m** and 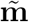 are both 3-vectors, and Ψ and 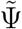 are both 3 × 3 positive semi-definite covariance matrices.

The full ALD covariance matrix (eq. 7), which multiplies together the diagonal matrices 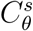 and 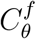 with a discrete Fourier domain operator in between, has the effect of simultaneously imposing locality in both space-time and frequency. It is identical to ASD under the setting that 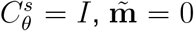, and 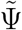 is diagonal [18]. Empirically, ALD has been shown to outperform both the ridge and ASD priors, as well as other sparsity-inducing priors, in applications to neural data in the early visual system [17].

## 4 A new prior for temporal Receptive Fields

Our first contribution is to propose a new prior for temporal receptive fields. The priors in ASD and ALD both assume a constant degree of smoothness across the receptive field. However, an assumption of stationary smoothness is less appropriate for temporal receptive fields, which are typically sharper at short time lags and smoother at longer time lags. In order to incorporate this variable form of smoothness, we introduce the Temporal Recency Determination (TRD) prior. This prior uses a smoothing covariance function that is stationary in log-scaled time, and which is therefore non-stationary in linear time.

The TRD covariance function can be described as a time-warped version of the ASD covariance. Specifically, the *i, j*’th element of the prior covariance is given by:

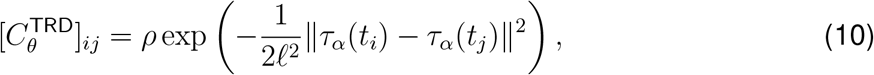

where *τ*_*α*_(*t*) is a nonlinear warping function:

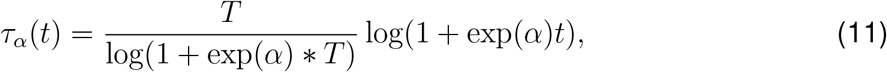

where *T* is the length of the temporal RF (in seconds), *t*_*i*_ is the time lag for temporal RF coefficient *i*, and *α* is a parameter determining how quickly the RF smoothness increases with time. Figure 2 shows an illustration of the TRD prior, alongside the other RF priors discussed in Sec. 3.

## 5 A probabilistic model for low-rank receptive fields

Our second contribution is a model and inference method for smooth, low-rank receptive field fields. As noted in Section 2.2, a low-rank parametrization for spatio-temporal receptive fields can offer a dramatic reduction in dimensionality without loss of accuracy. In particular, a rank *r* spatio-temporal filter requires only 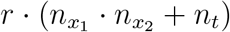 coefficients, versus 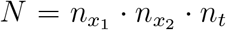 for a full-rank filter.

To place our model on a solid probabilistic footing, we place independent zero-mean Gaussian priors over the *r* temporal (1D) and *r* spatial (2D) components of the receptive field (see Fig. 3A). If we use a TRD prior for the temporal components and an ALD prior for the spatial components, then the prior over the *i*’th spatial and temporal receptive field components can be written:

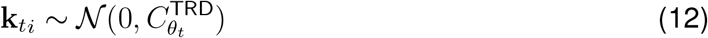

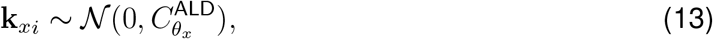

where 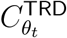 denotes the TRD covariance (eq. 10), 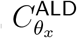 sis the ALD covariance (eq. 7), which we apply here to a 2D spatial receptive field. Although we selected these covariance functions because of their suitability for the structure of neural receptive fields, our modeling framework is general and could easily accommodate other choices.

**Figure 3:**
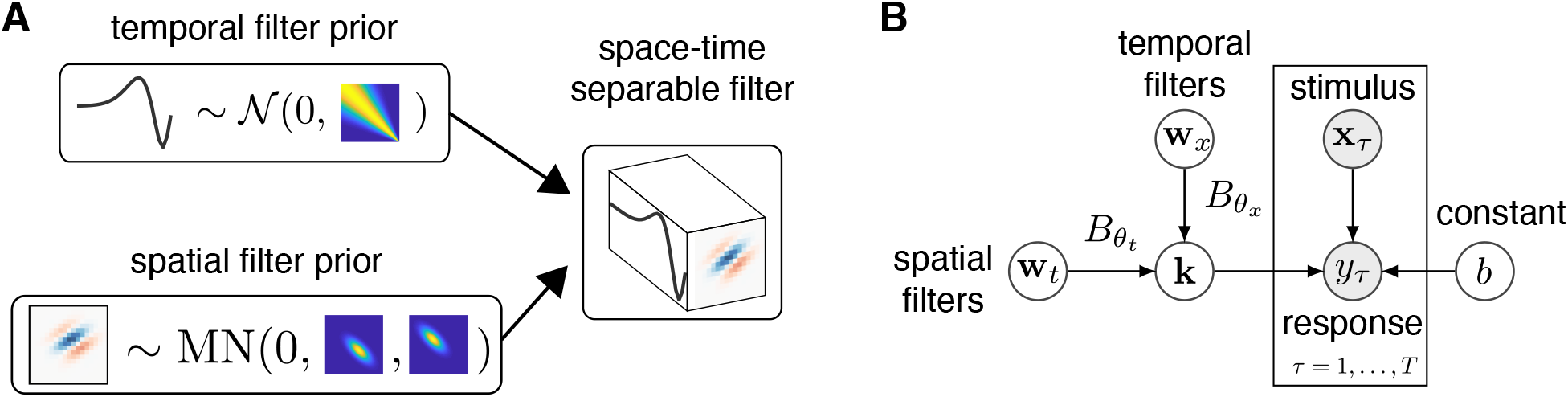
Graphical model representation of the low-rank STRF model. (**A**) We combine a multivariate normal ‘temporal relevance determination’ (TRD) prior over temporal components (above) and a matrix normal prior over spatial components (below), where row and column covariances have an ‘automatic locality determination’ (ALD) parametrization, to obtain a prior over each rank-1 component or space-time separable filter making up a low-rank filter. (**B**) The graphical model for the full model can be represented in terms of linear transformations 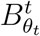 and 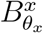, which parametrize the transformation from whitened temporal and spatial receptive fields **w**_*t*_ and **w**_*x*_, which are combined to form the STRF **k**. The linear transformations depend on hyperparameters *θ*_*t*_ and *θ*_*x*_. The spatial and temporal components are combined to form a low-rank STRF **k**, which acts to integrate a stimulus **x**_*τ*_ over space and time to generate a neural response *y*_*τ*_, with constant offset *b*.

Under this modeling choice, the full receptive field 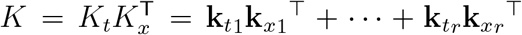 represents the product of two Gaussian random variables. This effective prior over *K* is thus non-Gaussian. This entails that posterior inference is not analytically tractable, due to the fact that the prior is not conjugate to the Gaussian likelihood of the encoding model (Eq. 1). We therefore develop a scalable variational inference algorithm for setting the hyperparameters *θ* governing the prior covariance matrices and inferring the receptive field coefficients *K*_*x*_ and *K*_*t*_, which we describe in sections 5.1-5.2. We introduce additional modelling choices that facilitate scalable estimation through a further reduction in the total number of receptive field coefficients in section 5.1, and describe the resulting algorithm in detail in section 5.2.

### 5.1 Scalability via sparse spectral representations of covariance matrices

Performing inference in the model we introduced in Section 5 still requires building and inverting large covariance matrices. To reduce the dimensionality of the spatial and temporal receptive field inference further and to achieve scalability to high-dimensional settings, we make use of sparse spectral representations of the prior covariance matrices.

This sparse spectral representation involves expressing the prior covariance of **k** as 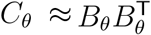. When *B*_*θ*_ contains the eigenvectors of *C*_*θ*_, scaled by their square-rooted associated eigenvalues, then this approximation is exact. However, we might be able to incur only a small approximation error by retaining only the leading eigenvectors of *C*_*θ*_ in *B*_*θ*_. For example, for larger lengthscale parameters (high degree of smoothness) the covariance matrix associated with the ASD covariance function has eigenvalues that quickly decay towards zero [28]. By truncating its eigenvalues near zero, it is possible to find a low-rank approximation to the covariance matrix *C*_*θ*_ that is exact to machine precision. For all choices of prior covariance matrices we discussed in section 3, it is possible to obtain an analytic expression for *B*_*θ*_ [18].

Given *B*_*θ*_, we can express the prior distribution over the receptive field in terms of a linear transformation of a multivariate standard Normal, or “whitened” random variable: **k** = *B*_*θ*_**w**, with **w** ~ 𝒩 (0, *I*). The resulting prior covariance of **k** is equal to 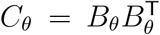. and thus unchanged from before. However, performing inference over **w** instead of **k** allows us to circumvent costly inversions of *C*_*θ*_. Furthermore, if *C*_*θ*_ is represented to a sufficient accuracy using fewer dimensions in *B*_*θ*_, then **w** will contain fewer dimensions than **k**, thus leading to additional improvements in scalability.

To incorporate low-rank receptive field structure into this approach, we define our model in the transformed, “whitened” space. The receptive field priors are described by multivariate standard Normal distributions over the whitened temporal receptive field **w**_*t*_ and whitened spatial receptive field **w**_*x*_:

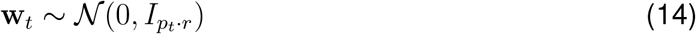

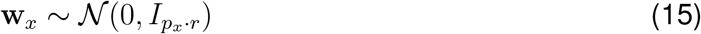

**w**_*t*_ and **w**_*x*_ are vectors containing the concatenated temporal and spatial components, respectively, and are of dimensions *p*_*t*_ · *r* and *p*_*x*_ · *r*, where *p*_*x,t*_ depend on the number of dimensions needed to approximate the respective covariance matrices to sufficient accuracy and *r* corresponds to the STRF rank. We define the matrix reshaping of these vectors as

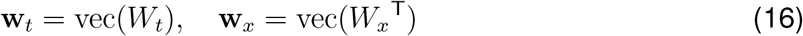

where 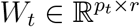 and 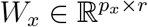. The full STRF can then be represented as

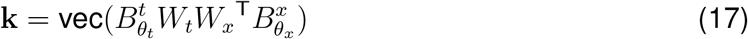

where the analytic decompositions of the prior covariances are denoted as 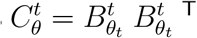 and 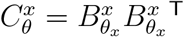 for the temporal and the spatial receptive field prior covariances, respectively.

Given this representation of the STRF, the rest of the model remains unchanged. This model description is illustrated as a graphical model in Figure 3B.

### 5.2 Variational inference and hyperparameter learning for low-rank STRFs (VLR)

Fitting our low-rank model to data involves performing posterior inference over the receptive field components **w**_*x*_ and **w**_*t*_, and learning the hyperparameters *θ* that determine the shape of the basis functions in *B*_*θ*_. To do this, we derive a variational optimization framework, similar to the classic Empricial Bayes approach reviewed in appendix A.2. We rely on a variational approach here, since the effective non-Gaussian prior over **k** makes obtaining an exact expression for the marginal log-likelihood intractable.

A lower bound to the marginal log-likelihood can be obtained by applying Jensen’s inequality

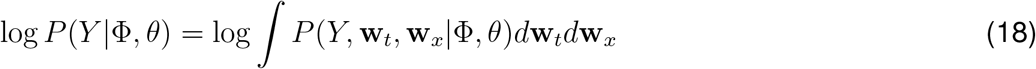

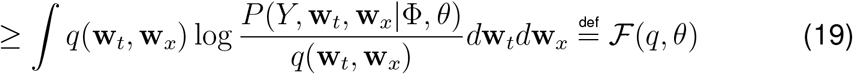

where *q*(**w**_*t*_, **w**_*x*_) is a distribution over the whitened temporal and spatial receptive field parameters **w**_*t*_ and **w**_*x*_. *ℱ* (*q, θ*) is often called the variational “free energy” or “evidence lower bound” (ELBO).

Instead of computing and optimising the model evidence directly, we can instead perform coordinate ascent, and alternatingly maximize ℱ with respect to the distribution *q*(**w**_*t*_, **w**_*x*_) and with respect to the parameters *θ*, as part of the Expectation Maximization (EM) [30] algorithm. The optimisation over *q*(**w**_*t*_, **w**_*x*_) can either be performed exactly (which amounts to setting *q* equal to the posterior joint distribution over **w**_*t*_ and **w**_*x*_), or under additional constraints on *q*, such as restricting it to a particular family of variational distribution.

The free energy can be written in two equivalent ways:

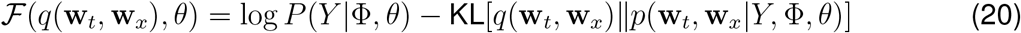

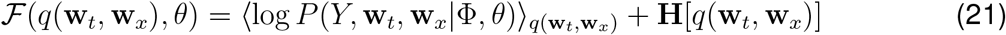

where **H**[*q*] is the entropy of *q*. The EM algorithm involves iteratively maximizing this lower bound with respect to a distribution over **w**_*t*_ and **w**_*x*_ (E-step) and with respect to the hyperparameters *θ* (M-step). From (20) it is apparent that the free energy is maximized when *q*(**w**_*t*_, **w**_*x*_) is equal to the posterior distribution

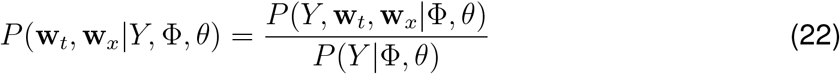

at which the Kullback-Leibler divergence in (20) vanishes and the free energy provides a tight lower bound to the marginal log-likelihood. Performing this computation exactly is intractable in our model. We therefore choose a variational approximation of the form

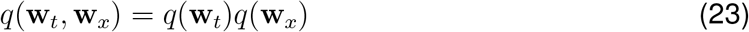

where we assume that the approximate posterior distribution factorizes over the spatial and temporal receptive field. We now seek to find the distribution *q*(**w**_*t*_, **w**_*x*_) that lies within the family of distributions that factorize over **w**_*t*_ and **w**_*x*_, and maximizes the free energy. Taking variational derivatives of the free energy, the variational updates for our approximating distributions are found to be of the general form

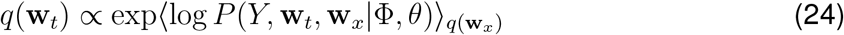

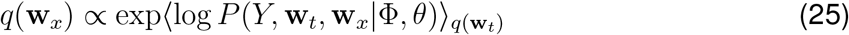

where angled brackets denote expectations. Evaluating the above in our model, we obtain multivariate Gaussian distributions of the form *q*(**w**_*t*_) = 𝒩 (**w**_*t*_ |*µ*_*t*_, Σ_*t*_) and *q*(**w**_*x*_) = 𝒩 (**w**_*x*_ |*µ*_*x*_, Σ_*x*_). The variational updates for the posterior means *µ*_*t*_, *µ*_*x*_, and covariances Σ_*t*_, Σ_*x*_ are available in closed form. Detailed derivations and the exact update equations are given in Appendix B.

The M-step in our variational EM algorithm involves maximizing the free energy with respect to the model parameters *b, σ* and hyperparameters *θ* = {*θ*_*t*_, *θ*_*x*_}, and requires solving

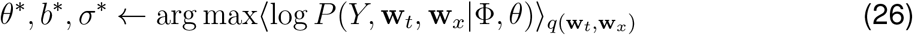

using gradient based optimization. We update the hyperparameters in 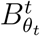 and 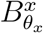, and the model parameters *b* and *σ* in separate conditional M-Steps. Performing conditional M-steps allows one to project the high-dimensional stimulus onto either the temporal or spatial basis and never requires building or inverting the full stimulus design matrix Φ. Thus, this strategy further exploits the lower-dimensional decomposition of the full receptive field and provides an efficient algorithm, even for high-dimensional data. Further details are provided in Appendix C.

## 6 Results

### 6.1 Data efficient estimation under correlated stimulus distributions

We first tested the VLR estimator using synthetic data. We simulated data from a linear-Gaussian encoding model with a rank-2 receptive field that was stimulated with correlated (exponentially filtered) Gaussian stimuli. To assess performance, we computed the estimate using different numbers of receptive field coefficients and different amounts of simulated training data. We compared VLR to the classic spike-triggered average (STA), and Bayesian ridge regression (RR). Figure 4A,B shows that the STA estimate suffers from severe bias due to the correlated stimulus distribution. The RR estimate provides an improvement in terms of the bias, but exhibits a high level of variability. Our proposed VLR estimator, in contrast, yields accurate estimates under correlated stimulus distributions, even when using only small amounts of training data.

**Figure 4:**
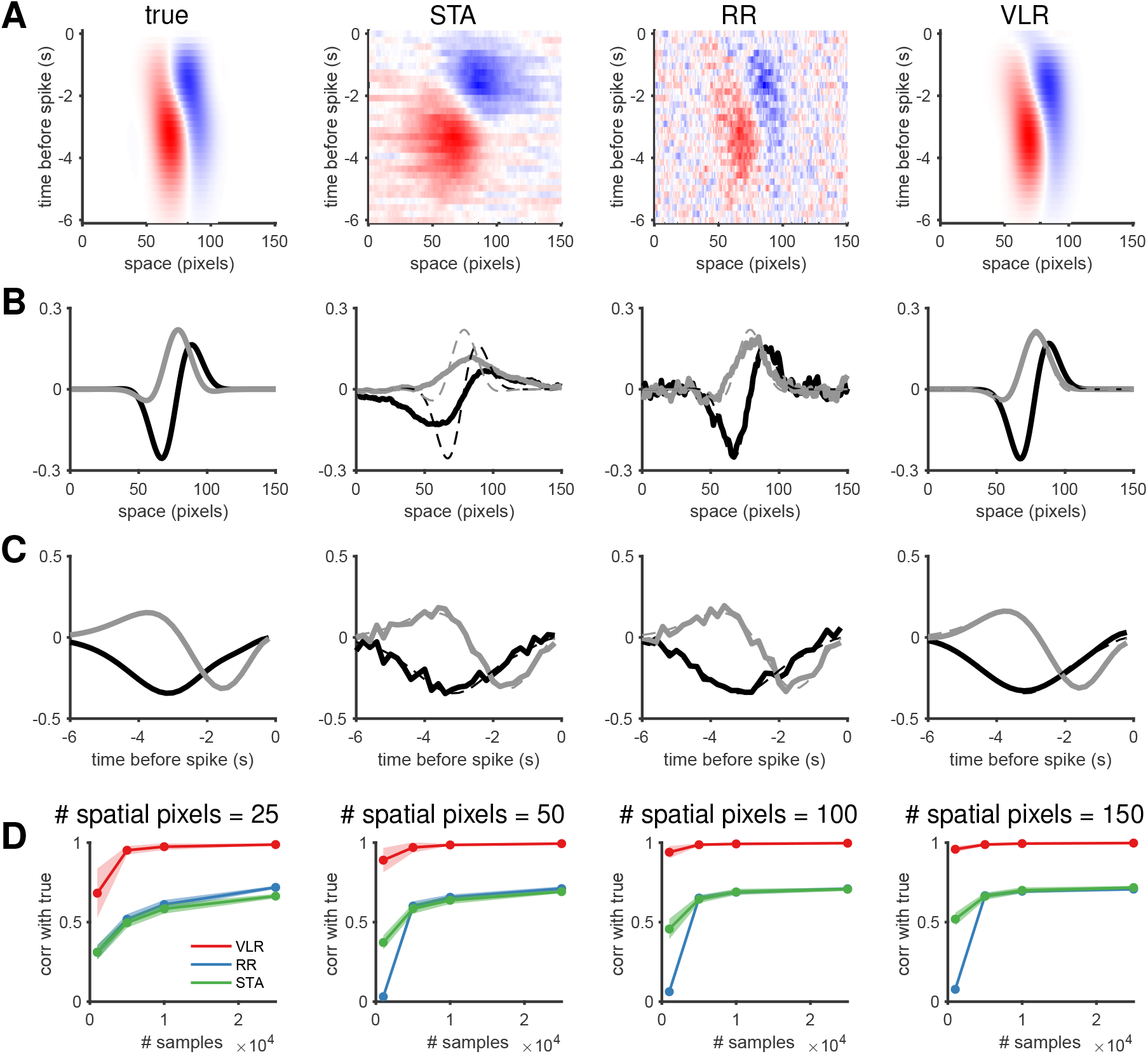
Synthetic data example. A: The true STRF that was used for generating synthetic responses and recovered estimates using STA, RR, and our VLR approach with a rank of 2, with ALD for the 2D spatial and TRD for the temporal component. Shown estimates are obtained using 1e^4^ training samples. B, C: Same as A but plotting the top two spatial (B) and temporal (C) components of the true and recovered STRF estimates. Dashed lines show the spatial and temporal components of the true STRF for reference. Spatial and temporal components of the STA and RR were not accessible by estimation, they were obtained by taking the leading left and right singular vectors of the STRF in matrix form and rescaling by the size of the true STRF. D: Correlation of the STRF estimate and the true STRF used to generate data, for different numbers of STRF coefficients and amounts of training data. Lines represent the average correlation across 20 repeated simulations with different random seeds; shaded regions represent ± 1 standard error.

### 6.2 Application to macaque V1 simple cells

Next, we examined recordings from V1 simple cells in response to random binary “bar” stimuli, previously published in [31]. The stimulus contained 12 to 24 1-dimensional spatial bars on each frame that were aligned with each cell’s preferred orientation. We used 16 time bins (160 ms) of stimulus history to define the temporal receptive field, so the entire STRF had between 12 × 16 and 24 × 16 total coefficients. We examined performance of estimator for different ranks and different amounts of training data, and evaluated the mean squared error on held-out test data. For comparison, we also computed the RR estimate and the Maximum Likelihood estimate (MLE), which in this setting corresponds to the ordinary least squares regression estimate (Fig. 5). We found that VLR outperformed the MLE and RR estimators, achieving lower prediction errors on held-out test data under varying amounts of training data (Figure 5C,F,I). These results illustrate the heterogeneity of STRF ranks in the V1 simple cell population, as the three example STRFs shown in Figure 5A,D,G achieved minimal prediction error for different choices of STRF rank (Figure 5B,E,H).

**Figure 5:**
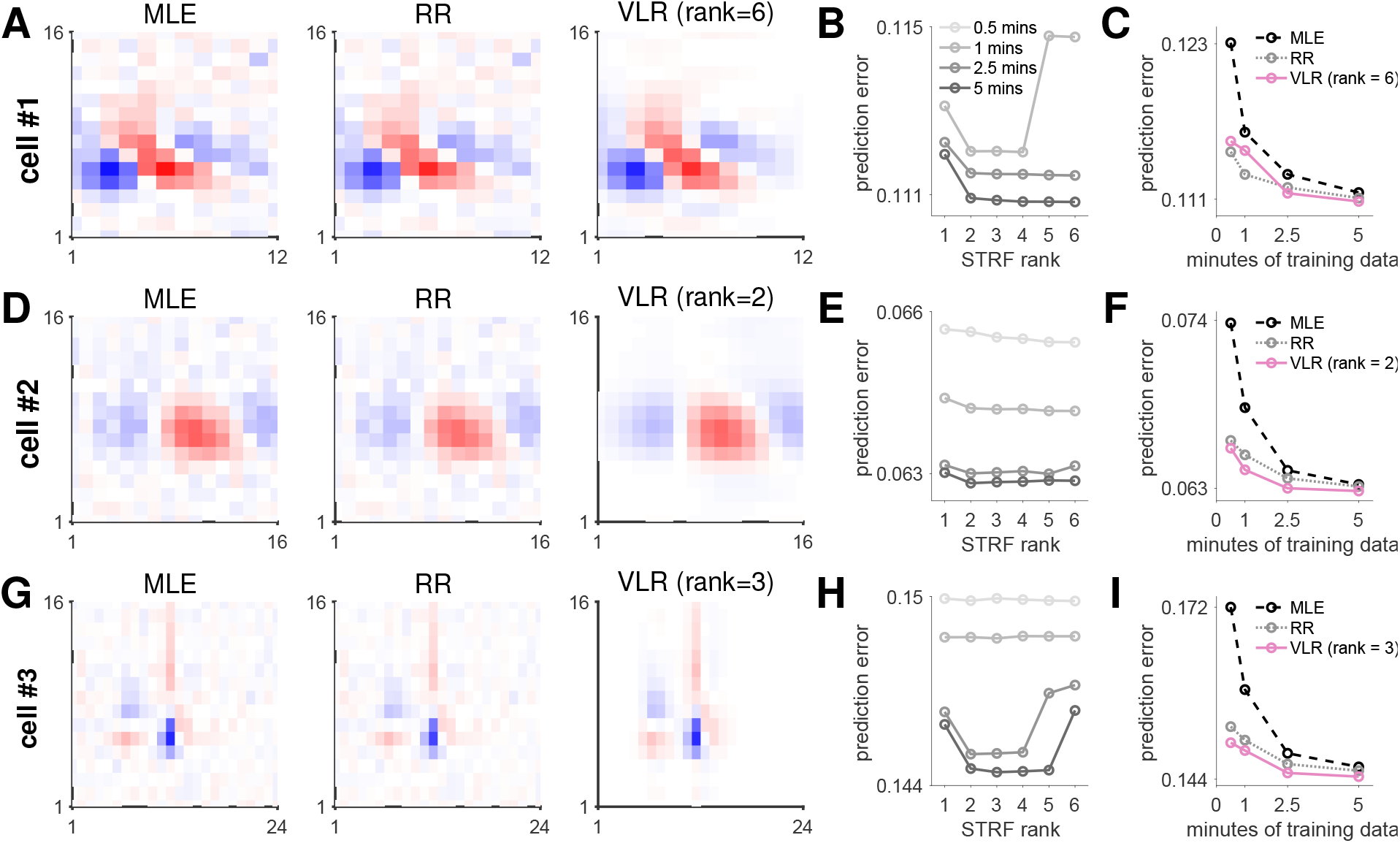
Application to V1 simple cells **A**,**D**,**G**: Example STRF estimates using MLE, RR, and our VLR approach. Shown estimates are obtained using 5 minutes of training data. **B**,**E**,**H**: Mean squared error on 10 minutes of held-out test data using different ranks and amounts of training data for the VLR approach. **C**,**F**,**I**: Mean squared error on 10 minutes of held-out test data when using different amounts of training data for MLE, RR, and an example rank for VLR.

### 6.3 Application to rat retinal ganglion cells

Previous work has identified multiple cell types of retinal ganglion cells (RGCs) [32]. Functional cell-type distinctions are usually based on receptive field properties such as ON/OFF regions, temporal filtering properties, or other structure that can be extracted from STRFs [33–35]. Other differences in response properties, such as STRF rank, may prove valuable for further distinguishing functional cell-types.

To further illustrate the power of the VLR estimator, we examined a dataset consisting of electrophysiological recordings from rat RGCs in response to binary white noise stimuli. We selected a region of 25 × 25 pixels in space based on the center of the RF (as estimated by the STA) from the 80 × 40 overall stimulus, and we considered 30 time lags (0.5 seconds) of stimulus history. This yielded a total of *N* = 18750 STRF coefficients. Standard regression estimators like the MLE require large amounts of training data for STRFs of this size. Bayesian estimators like ASD and ALD are computationally intractable in high-dimensional settings due to necessity of storing and inverting matrices of size *N* ^2^. Thus, we compare the VLR estimator with the STA, which can readily be computed even for large STRF sizes such as those considered here.

Figure 6 shows the performance of STA and VLR for different receptive field ranks. VLR achieved lower prediction errors on held-out test data, even when using only small amounts of training data. Figure 6A shows that VLR outperformed the STA estimate computed on 83 minutes of training data using as little as 4.2 minutes of training data, and under all assumed STRF ranks (Figure 6B). Comparing the STA and VLR estimates in the top panels of Figures 6D,E, VLR managed to more successfully extract signal from the data and reduce speckle noise in the estimate. The temporal and spatial components of the STRF can be extracted as the leading left and right singular vectors of the matrix unfolding of the STRF tensor, each weighted by the square-root of the associated sigular value. The STA estimate was dominated by noise and provided much poorer estimates of the spatial and temporal components of the STRF, as demonstrated in the bottom panels of Figure 6D,E. The second spatial component of the STA estimate is dominated almost exclusively by speckle noise (6D, lower left), while it is structured in the VLR estimate (6E, lower left). The rank of the receptive field indicates how many rank-one space-time components are required to reconstruct the STRF. The singular values of the STRF indicate the associated weight of each rank-one space-time component in this reconstruction. As the assumed rank of the VLR estimate grows, our model has the capacity to fit more complex structure in the STRF. Figure 6C shows the singular values of the STA and and VLR STRF estimates under different assumed ranks. As the assumed rank of the VLR estimate grows, the associated singular values decay to zero. This demonstrates that the VLR estimator is able to successfully prune out space-time components that do not reflect signal in the training data, and thus prevents overfitting to noise. This feature can be attributed to the non-Gaussian prior distribution over the effective STRF weights of our model, and is a favorable regularization property. As a result, the prediction error plateaus after a sufficient rank has been reached, demonstrating that even estimates obtained under a higher assumed rank generalize well to unseen data (Figure 6B).

**Figure 6:**
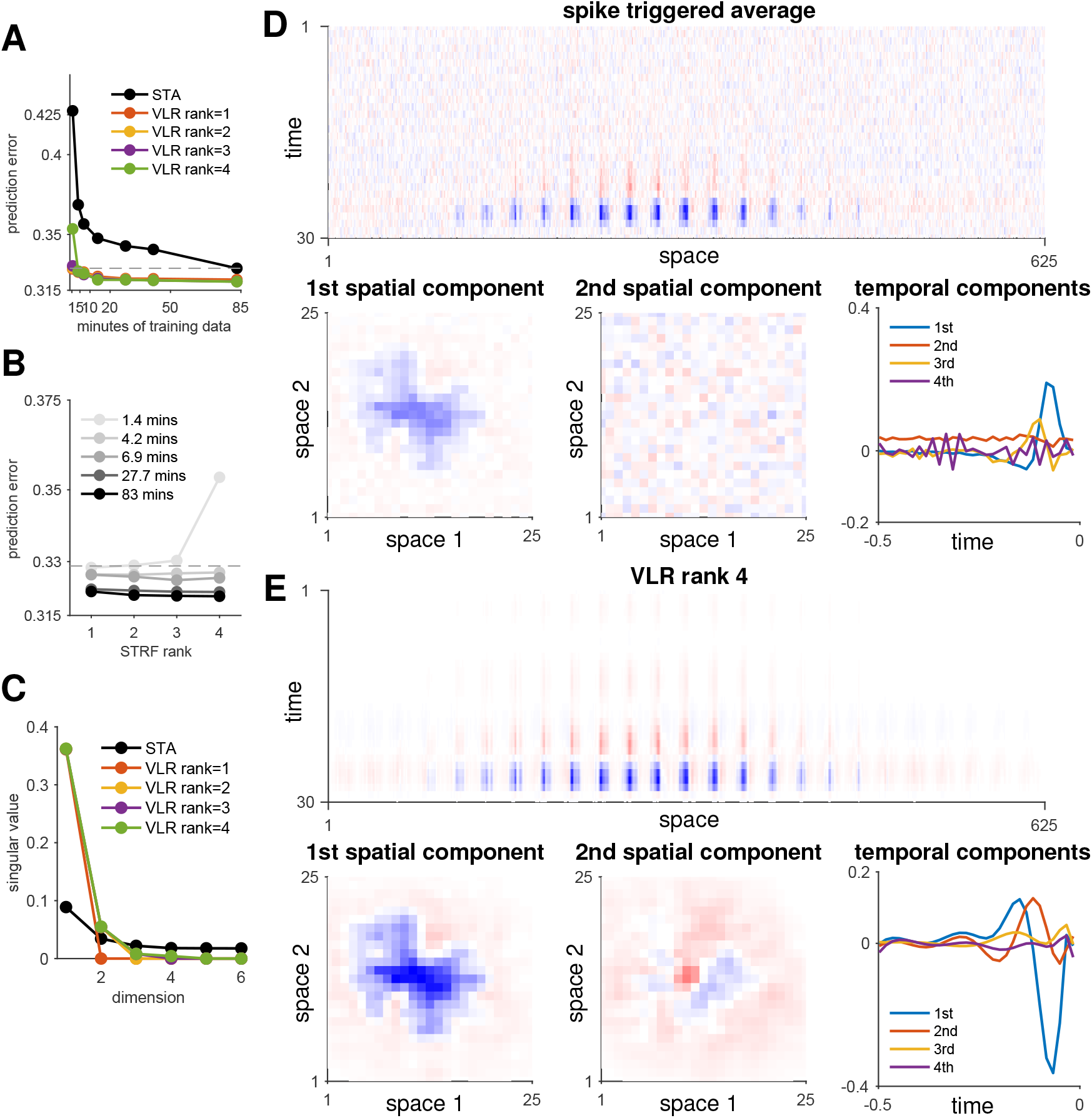
Application to a rat retinal ganglion cell (RGC). **A**: Mean squared error on 27.66 minutes of held-out test data using different ranks and amounts of training data. The dashed grey line indicates the prediction error achieved by the STA using the maximum amount of training data, showing that estimates computed under the VLR approach achieve similar performance using as little as 4.2 minutes of training data. **B**: Same as **A** but plotting the prediction error against ranks for the VLR approach. **C**: The top singular values computed on the STRF estimate using STA or the VLR approach under different assumed ranks and using the maximum amount (83mins) of training data. **D**: STRF estimate computed using STA on 13.83 minutes of training data together with the top two spatial and top four temporal components, determined as the leading left and right singular vectors of the STRF, scaled by the square root of the corresponding singular value. **E**: Same as **D** but using the VLR approach with an assumed rank of 4.

Lastly, in Figure 7 we show summary statistics across a population of 15 RGCs, which can be grouped into three cell-types depending on STRF properties such as ON/OFF regions or STRF shape. Figure 7A shows that VLR robustly outperforms the STA, while Figure 7B shows that STRF ranks are diverse across the assigned cell-types, adding to previous reports on diverse response properties within ON/OFF pathways in mammalian RGCs [36]. Figure 7C shows the leading spatial component of the inferred STRFs across different cells and cell-types. Further examples along with the temporal receptive field components are shown in Supplementary Figure 1.

**Figure 7:**
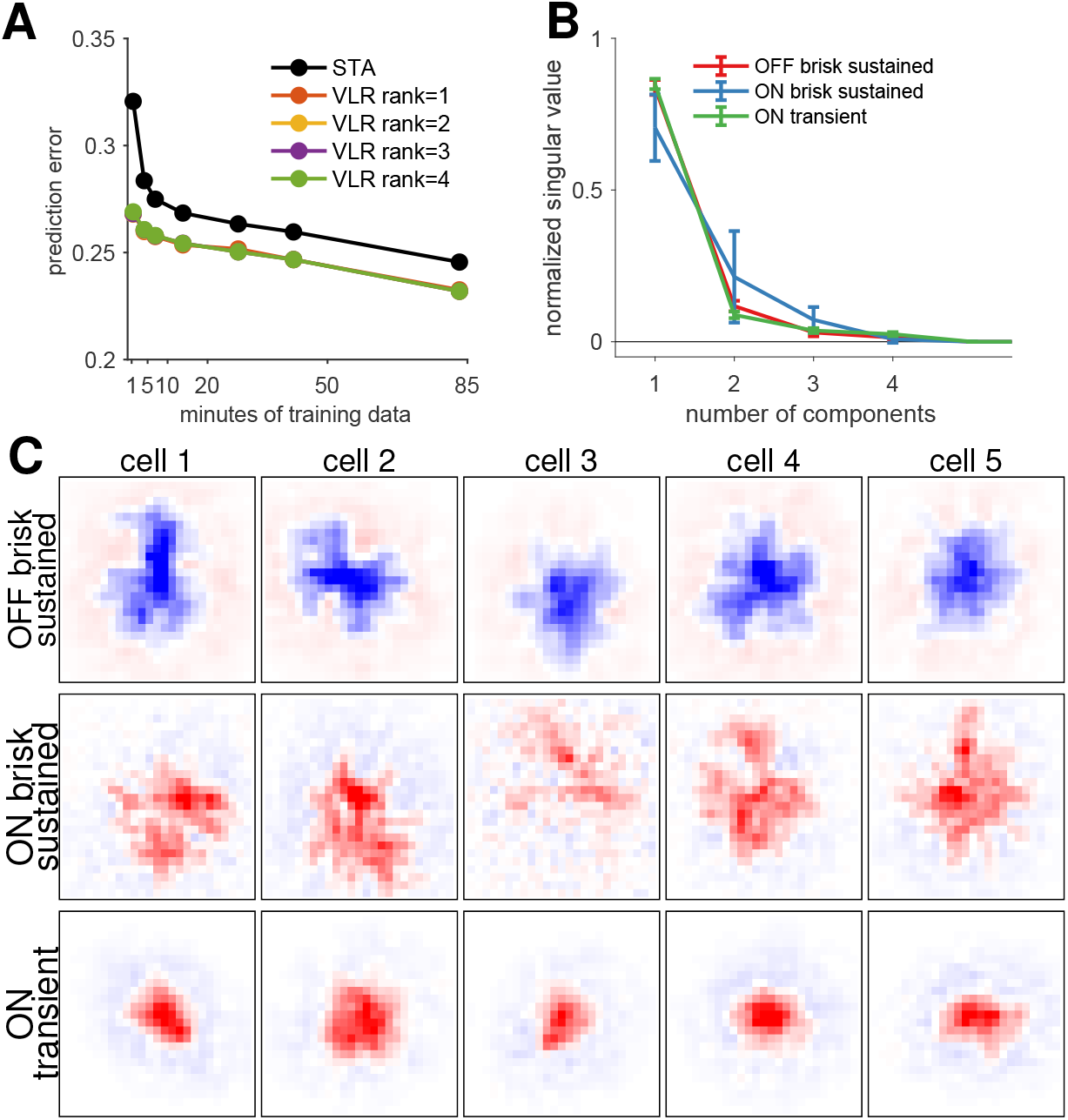
Application to a population of rat retinal ganglion cells. **A**: Mean squared error on 27.66 minutes of held-out test data using different ranks and amounts of training data, averaged across 15 cells. **B**: Mean and *±* one standard deviation of the singular values for the rank 4 VLR estimate, separated by cell-type. Singular values are normalized to sum to one. Cell-types have been classified by hand by the experimenters based on STRF properties. **C**: Leading spatial STRF component for five example cells of each of three different cell-types.

## 7 Discussion

In this paper, we have introduced a novel method for inferring low-rank STRFs from neural recordings, and have shown a substantial improvement over previous methods for STRF estimation. Previous approaches like the STA have low computational complexity and are hence applicable to STRFs with large sizes, but provide biased estimates under correlated stimulus distributions and classically require large amounts of training data to arrive at accurate estimates. Previous Bayesian approaches have been shown to be more data efficient and can provide more accurate STRF estimates, even under correlated stimulus distributions. However, they require matrix inversions that are computationally costly. Thus, applying Bayesian approaches to large stimulus ensembles remains challenging. Furthermore, all of these approaches parameterize the full STRF and do not explicitly make use of potential low-rank structure in the STRF. These methods are therefore general, but require estimation of large numbers of parameters which prohibits their application to large-scale problems.

The VLR estimator we have introduced in this paper addresses limitations of previous approaches to STRF estimation in linear encoding models. We have shown that our method provides accurate estimates in settings when stimuli are correlated, the receptive field dimensionality is high, or training data is limited. We developed a new prior covariance function that captures the non-stationary smoothness that is typical of temporal receptive fields. In combination with previous priors capturing localized smooth structure for spatial receptive fields, our probabilistic model provided a powerful framework for low-rank STRF estimation. While we have focused on these modeling choices in this paper, our modeling framework is general and can easily accommodate other choices of prior covariance functions.

The ability to accurately estimate STRFs from limited amounts of data will be important for quantifying changes of receptive field properties over time, or during learning. Furthermore, it will open up possibilities for closed-loop experiments, where stimuli are selected actively on each frame to reduce uncertainty about the STRF. Furthermore, being able to use high-dimensional stimuli and correlated stimulus distributions will be important for studying population responses. In this setting, VLR will be useful for estimating the receptive fields of many cells in parallel. Finally, VLR allows for cross-validated estimates of the STRF rank, which may prove important for quantifying single-cell response properties and categorize cells into specific functional classes.

While we have presented our method in the context of Gaussian linear encoding models, it is also possible to extend the variational inference algorithm we presented here to non-Gaussian observation models in a generalized linear encoding model (GLM). Here, the ability to perform closed form updates for the posterior means and covariances will be lost and will instead require direct optimization of the variational free energy. Previous aproaches have made use of basis functions to improve the scalability of RF estimation in Poisson GLMs [20]. However, this typically requires setting parameters like the number, location and shape of basis function by hand. Recent work has started to consider circumventing this cumbersome by-hand selection of basis functions by using sparse variational Gaussian process approximations instead [37]. This approach relies on inducing point methods which recently gained popularity in improving the scalability of Gaussian Process methods more generally [38, 39]. The framework we presented here is closely related to the basis function approach for GLMs and can be used to determine parameters such as the number, location and shape of basis functions automatically. The improvements in scalability we achieve due to whitened representations of the receptive field also apply in the context of GLMs. Ultimately, algorithms that can automatically determine such hyperparameters and do not require user-dependent input will contribute to the robustness and reproducability of modeling results.

## Code availability

We provide `Matlab code` for fitting low-rank STRFs under flexible choices of prior covariance functions at https://github.com/pillowlab/VLR-STRF.

## Author Contributions

LD and JWP conceived of this project. LD developed the presented methods and performed analyses under supervision of JWP. KMR and GDF collected the retinal ganglion cell experimental data and assisted in the retinal ganglion cell analyses. LD and JWP wrote the manuscript with input from KMR and GDF.

## Acknowledgements

We would like to thank Sneha Ravi for sharing the RGC data, and Nicole Rust and Tony Movshon for providing access to V1 simple cell data. LD was supported by the Gatsby Charitable Foundation. JWP was supported by grants from the McKnight Foundation, the Simons Collaboration on the Global Brain (SCGB AWD543027), the NIH BRAIN initiative (NS104899 and R01EB026946), and a U19 NIH-NINDS BRAIN Initiative Award (5U19NS104648).

## A Bayesian receptive field estimation

Here we review the standard approaches for estimating receptive fields from data under the linear-Gaussian encoding model (Sec. 2) using Gaussian priors (Sec. 3).

### A.1 Maximum a posteriori (MAP) estimation

Given a fixed setting of the hyperparameters *θ*, it is straightforward to compute the *maximum a posteriori* (MAP) estimate, which is simply the maximum of the posterior distribution over **k** given the data. The posterior can be computed using Bayes’ rule:

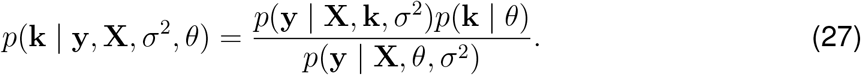

The numerator consists of the likelihood times the prior, where the likelihood term comes from the Gaussian encoding model,

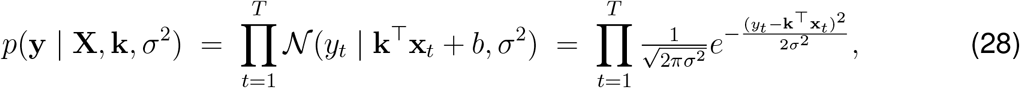

and the prior is

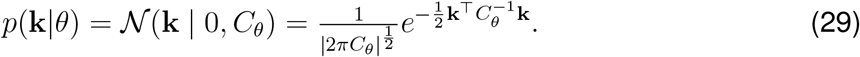

The denominator term, *p*(**y** | **X**, *θ*), is known as the *evidence* or *marginal likelihood*, and represents a normalizing constant that we can ignore when computing the MAP estimate.

In this setting, where the likelihood and prior are both Gaussian in **k**, the posterior is also Gaussian. The posterior mean, which is also the MAP estimate, has a closed-form solution:

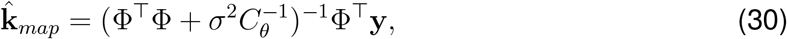

where **y** denotes the vector of responses, and Φ is the design matrix, which contains the corresponding stimulus vectors along its rows:

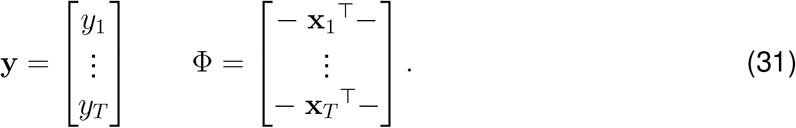

If needed, the posterior covariance is equal to

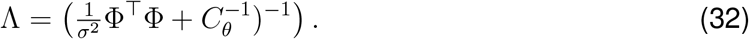

### A.2 Evidence optimization and empirical Bayes

A critical question we have not yet discussed is how to set the noise variance *σ*^2^ and the hyperparameters *θ* governing the prior. A common approach for selecting hyperparameter values is cross-validation over a grid of candidate values. However, this approach is unwieldy for models with more than one or two hyperparameters.

A popular alternative in Bayesian settings is evidence optimization, also known as type-II maximum likelihood estimation [17, 25, 40, 41]. The idea is to compute maximum likelihood estimates for *σ*^2^ and *θ* using the marginal likelihood or *evidence*, which is obtained by marginalizing over the model parameters **k**. Incidentally, the marginal likelihood is also the denominator in Bayes’ rule (eq. 27). In the current setting, the marginal likelihood is given by:

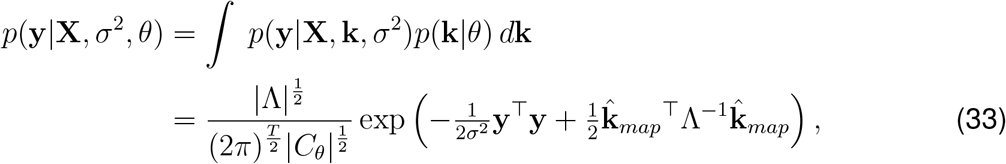

where 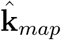 and Λ are the posterior mean and covariance defined above.

In practice, one performs inference for *σ*^2^ and *θ* by numerically optimizing the log of the marginal likelihood:

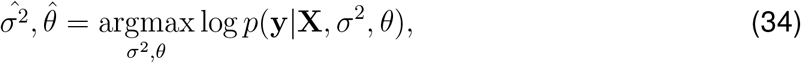

although there are also fixed-point methods available for the ridge regression case [17, 40]. Once this numerical optimization is complete, we can compute the MAP estimate for the receptive field **k** conditioned on these point estimates. This two-step estimation procedure (evidence optimization followed by MAP estimation) is known as *empirical Bayes*:

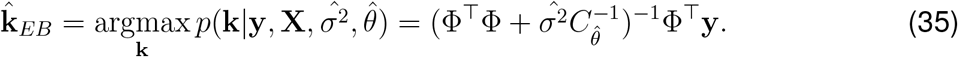

This approach has been shown to obtain substantially improved receptive field estimates in settings with limited data and correlated stimulus distributions, particularly using ASD or ALD priors [17, 25]. However, computing the evidence (eq. 33) or even the simple MAP estimate (eq. 30) requires storing and inverting a matrix of size *N × N*, where 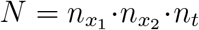 is the total number of coefficients in **k**. The storage costs thus scale as *O*(*N* ^2^), while the computational costs scale as *O*(*N* ^3^), cubically in the number of receptive field coefficients. This severely limits the feasibility of MAP and empirical Bayesian estimators in high-dimensional settings.

## B Variational updates for low-rank STRF inference

The variational updates require computing

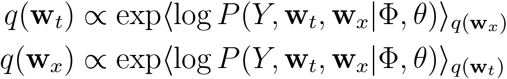

Both of these distribution will turn out to be Gaussians, such that the variational update, or E-step, reduces to computing means and covariances that fully specify the distributions *q*(**w**_*t*_) = 𝒩 (**w**_*t*_|*µ*_*t*_, Σ_*t*_) and *q*(**w**_*x*_) = 𝒩 (**w**_*x*_|*µ*_*x*_, Σ_*x*_).

To find the variational update for *q*(**w**_*t*_), we evaluate the expected log-joint distribution of the response and receptive field with respect to *q*(**w**_*x*_). The log-joint distribution can be expressed as

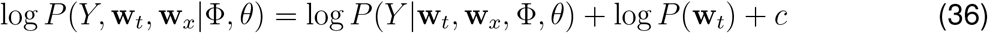

where *c* absorbs all terms that are constant with respect to **w**_*t*_. The log-likelihood of the response can be expressed as

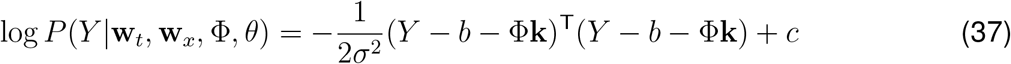

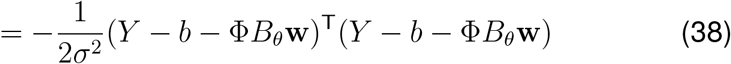

Using properties of the Kronecker product, we can note that **w** can be re-written in two ways:

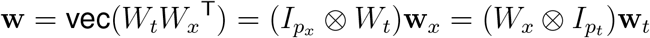

with **w**_*t*_ = vec(*W*_*t*_) and **w**_*x*_ = vec(*W*_*x*_^T^). Substituting this into the expression for the log-joint distribution and taking expectations (denoted by angled brackets) we have

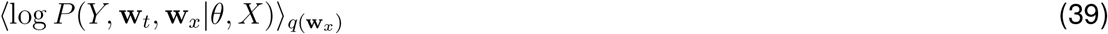

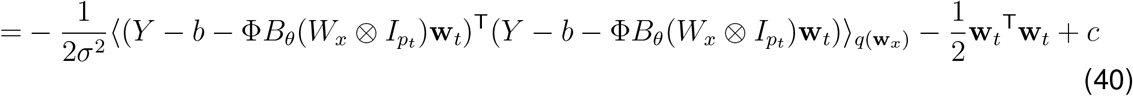

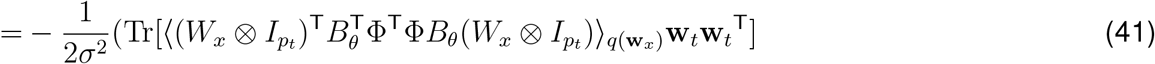

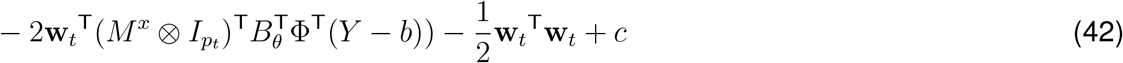

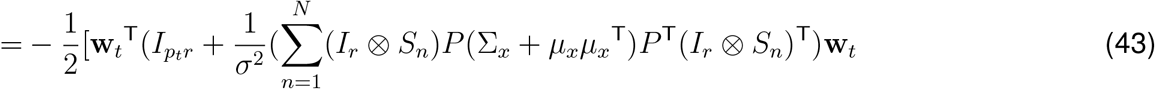

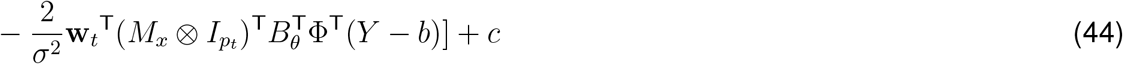

where *µ*_*x*_ = vec(*M*^*x*T^) = vec(⟨*W*_*x*_^T^⟩_*q*(**w**)_), *P* is a commutation matrix [42] such that vec(*M*_*x*_) = *P* vec(*M*^*x*T^) = *Pµ*_*x*_, and *S*_*n*_ = reshape 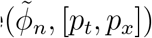 and 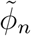 is the *n*-th row of Φ*B*_*θ*_. The above can be brought into a multivariate Gaussian form by completing the square, yielding the variational updates:

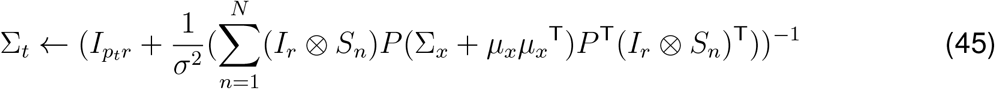

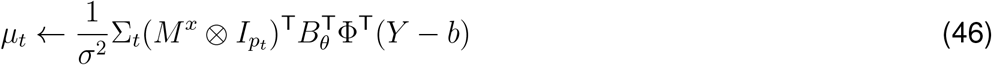

Similarly, to evaluate the expectation with respect to *q*(**w**_*t*_):

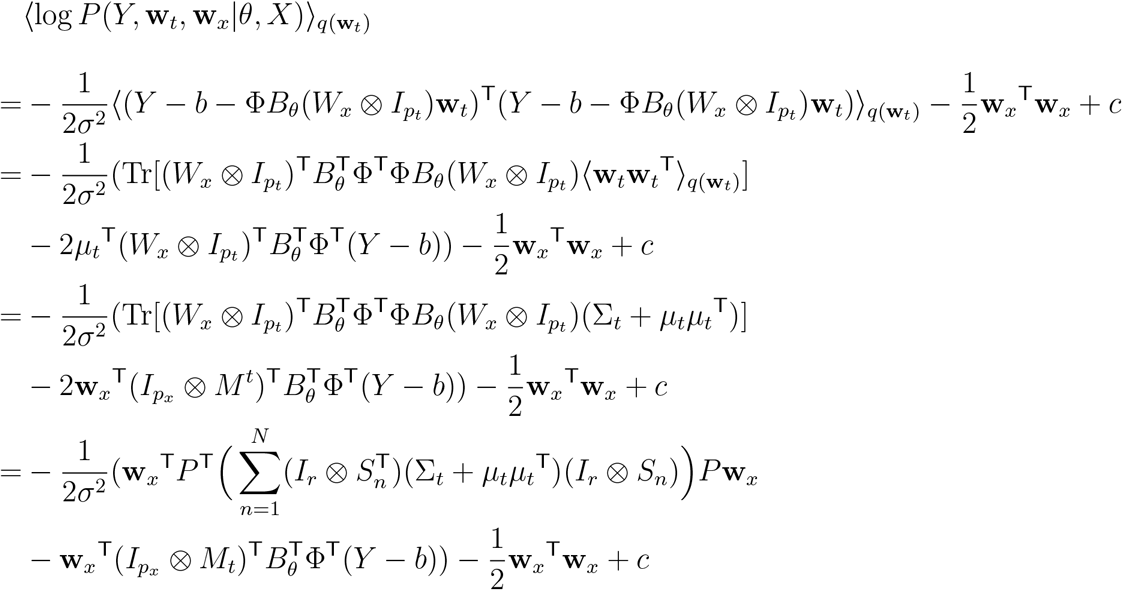

where *µ*_*t*_ = vec(*M*^*t*^) = vec(⟨*W*_*t*_ ⟩_*q*(**w**)_), and *S*_*n*_ and *P* are the same as before.

We can again complete the square to obtain a Gaussian form in **w**_*x*_, leading to the variational updates

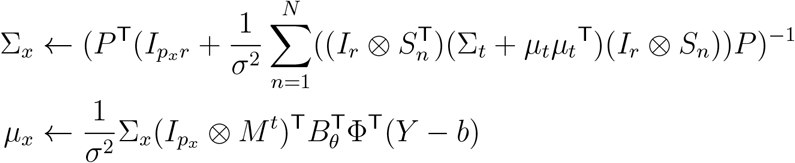

## C M-Step updates for low-rank STRF inference

The M-step updatesfor the hyperparameters *θ* = {*θ*_*x*_, *θ*_*t*_} can be carried out most efficiently by performing conditional M-steps for either the spatial or temporal parameters, while holding the other set of parameters fixed.

Dropping all terms that do not depend on the hyperparamaters *θ* from the free energy:

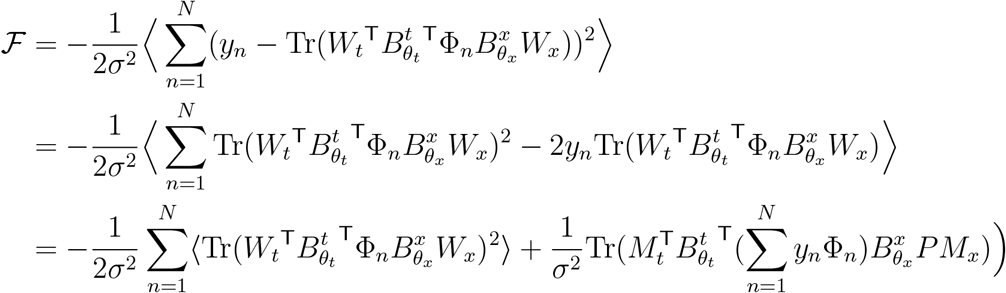

Here Φ_*n*_ is the *n*th row of Φ (which is equal to **x**_*n*_), reshaped to have the same size as the STRF.

To evaluate the first term efficiently, it can be rewritten as follows:

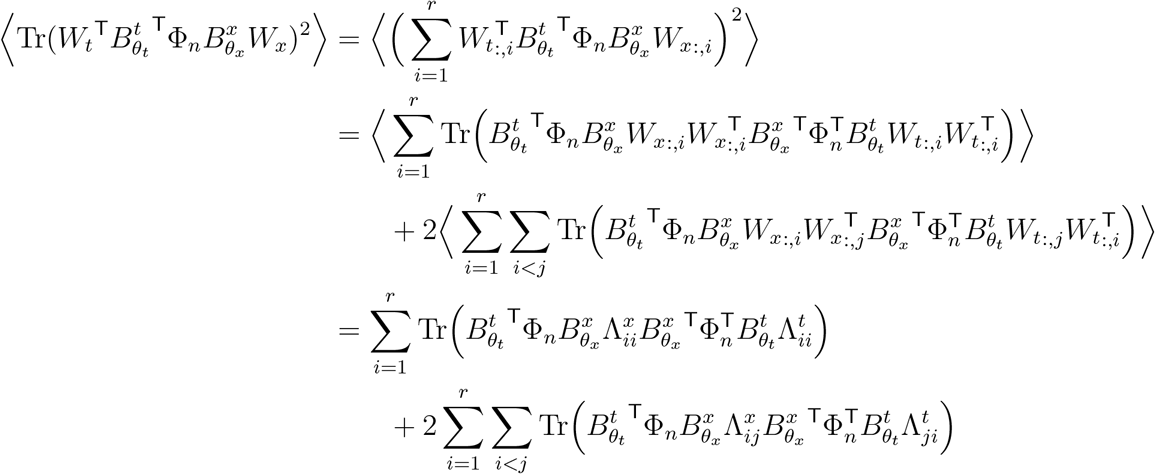

Here, the blocks of the second moment of the vectorized STRF are expressed as

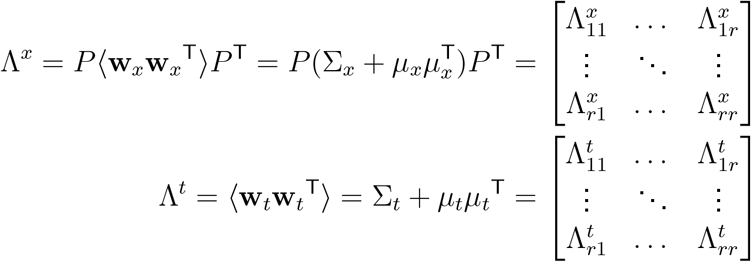

It is apparent from the above expression that the conditional M-steps will rely on statistics of the stimulus projected onto the basis we are not optimizing over. In order to evaluate each M-step update as efficiently as possible, these statistics can be pre-computed and stored throughout the M-step update. Below, we provide further details for each conditional M-step.

### C.1 Spatial parameter updates

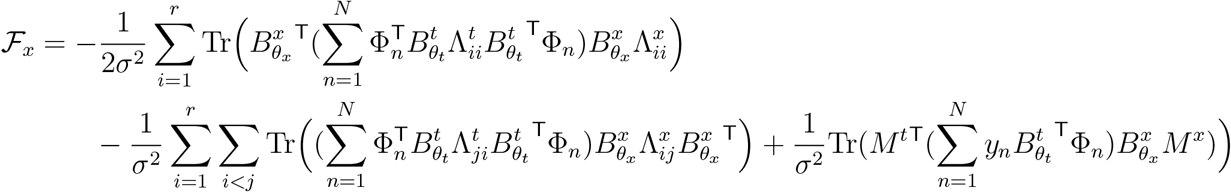

Now, before computing the parameter updates for the spatial parameters, we can pre-compute:

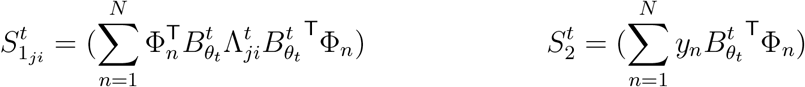

This involves 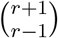 matrices of size *p*_*x*_ *× p*_*x*_. While building this matrix is expensive, the size of *p*_*x*_ is pruned using ALD. This cost is only incurred once before performing repeated updates to the spatial hyperparameters.

### C.2 Temporal parameter updates

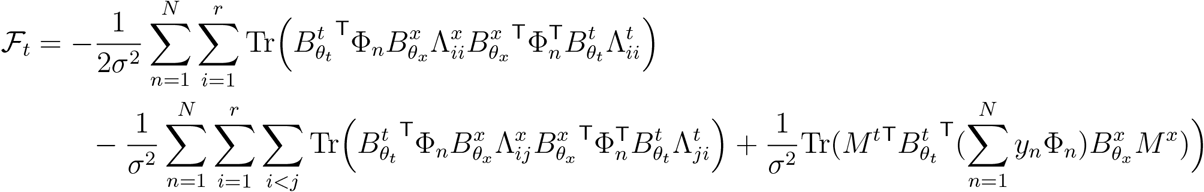

Now, before computing the parameter updates for the spatial parameters, we can pre-compute:

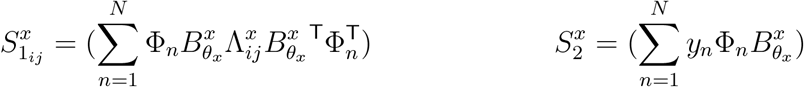

Analogously to the spatial parameter updates, this involves pre-computing and storing 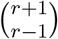 matrices of size *p*_*t*_*× p*_*t*_.

**Supplementary Figure 1.**
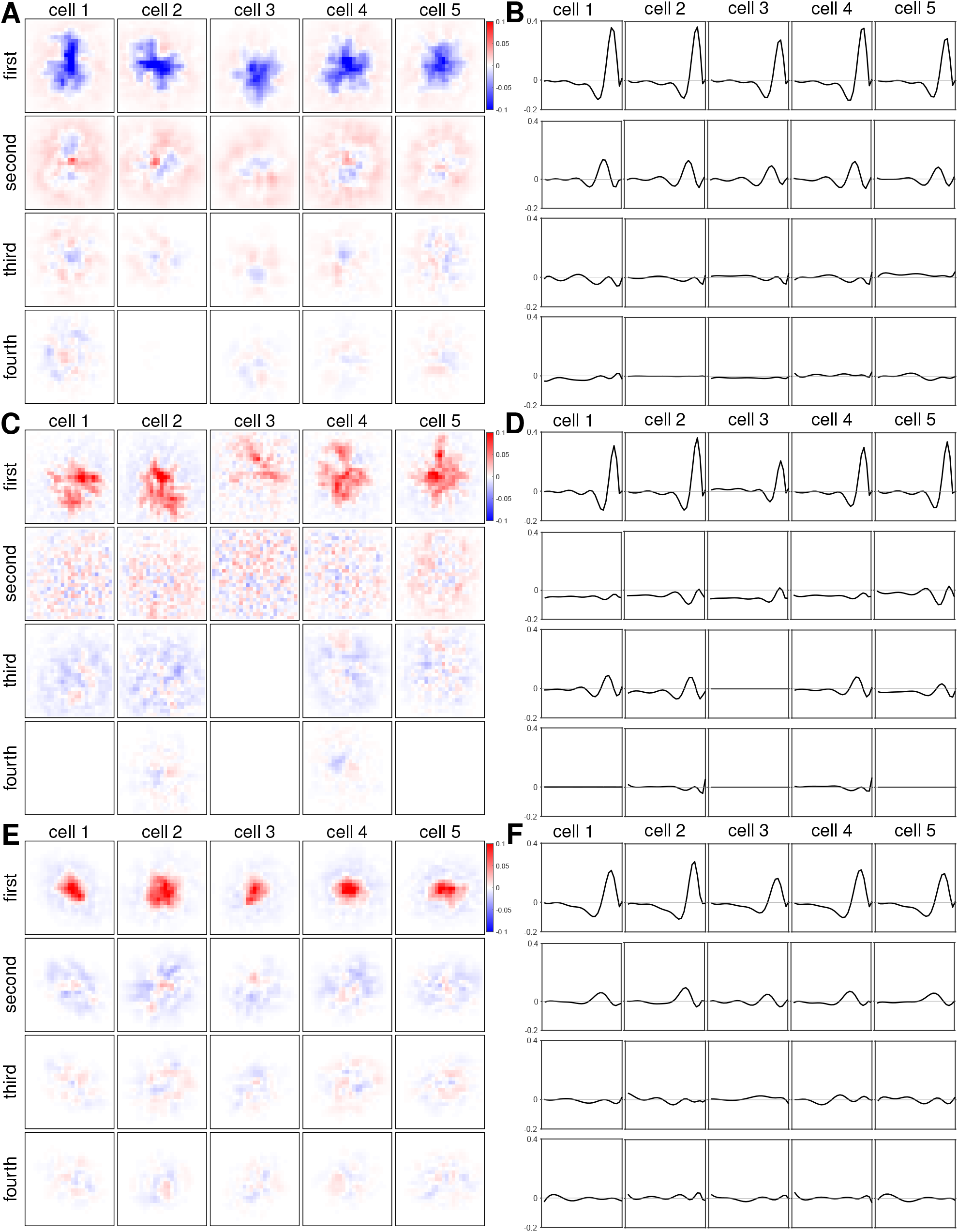
Inferred spatial and temporal components using VLR for different cell types and example cells. Each panel shows the four spatial (**A**,**C**,**E**) and temporal (**B**,**D**,**F**) low-rank receptive field components for five example cells that have been classified as *OFF brisk sustained* (**A**,**B**), *ON brisk sustained* (**C**,**D**), or *ON transient* (**E**,**F**).

## Notes

### Competing Interest Statement

The authors have declared no competing interest.

